# Creativity in Verbal Associations is Linked to Semantic Control

**DOI:** 10.1101/2022.02.08.479385

**Authors:** Katya Krieger-Redwood, Anna Steward, Zhiyao Gao, Xiuyi Wang, Ajay Halai, Jonathan Smallwood, Elizabeth Jefferies

**Author notes:** Corresponding author, Department of Psychology, The University of York, Heslington, York, PS/E008, YO10 5DD, UK.

## Abstract

While memory is known to play a key role in creativity, previous studies have not isolated the critical component processes and networks. We asked participants to generate links between words that ranged from strongly related to completely unrelated in long-term memory, delineating the neurocognitive processes that underpin more unique versus stereotypical patterns of retrieval. Less creative responses to strongly associated word pairs were associated with greater engagement of episodic memory: in highly familiar situations, semantic and episodic stores converge on the same information. This pattern of retrieval was associated with greater engagement of core default mode network. In contrast, more creative responses to weakly associated word pairs were associated with the controlled retrieval of less dominant semantic information and greater recruitment of the semantic control network, which overlaps with the dorsomedial subsystem of default mode network. Consequently, although both creative/controlled and stereotypical/more automatic patterns of retrieval are associated with activation within default mode network, these processes show little overlap in activation. These findings show that creativity emerges from controlled aspects of semantic cognition.

---

Creativity and communication depend on our capacity to deploy information from memory in a flexible way. As an illustration, we can generate an association between *any* two words (even unrelated items) by identifying a specific context in which they interact or co-occur (e.g. we can associate the words MELON and BOOKCASE by thinking about cookery books); this behaviour is highly creative since there is no obvious way in which these words are linked. Creativity is assumed to reflect the ability to generate unusual patterns of retrieval from memory – including from the semantic store (encompassing the meanings of words and objects; Abraham & Bubic, 2015; Chen et al., 2015; Kenett, 2018; Kenett & Faust, 2019; Liu et al., 2020; Mednick, 1962) and/or from episodic memory (which represents our individual experiences; Addis et al., 2016; Beaty et al., 2016; Benedek & Fink, 2019; Madore et al., 2015; 2016a; 2016b). Previous research has shown that executive and default mode networks are recruited during creative thought (Beaty et al., 2015; 2016; 2014; 2019), yet the component processes reflected by these network interactions remain unclear. Neuroscientific studies of memory have revealed distinct neural networks that are engaged during controlled as opposed to automatic patterns of retrieval from both semantic and episodic memory (Barredo et al., 2015; Davey et al., 2016; Kim, 2016; Vatansever et al., 2021; Whitney et al., 2009), yet these studies typically only examined judgements about pre-linked words, minimising the contribution of creativity. This study therefore investigated neural recruitment as participants formed links between words that varied in their degree of association along a continuum from strongly related (lowest creativity) to unrelated (highest creativity), linking neural activation and behavioural performance to distinct aspects of memory.

Previous studies have found that the efficient activation of broader conceptual information increases the likelihood of creating unique conceptual combinations (Benedek et al., 2017; Kenett et al., 2014; Kenett et al., 2016; Kenett & Faust, 2019). This pattern of retrieval may be connected to controlled semantic cognition, as opposed to the retrieval of conceptual knowledge that comes unbidden to mind, since sematic control processes are thought to be key to the retrieval of weak, ambiguous or non-dominant aspects of knowledge (Badre & Wagner, 2007; Hoffman et al., 2010; Jefferies, 2013; Krieger-Redwood et al., 2015). In these circumstances, a left-lateralised semantic control network (SCN) is strongly activated: this includes left inferior frontal gyrus (IFG), posterior middle temporal gyrus and dorsomedial prefrontal cortex (Jackson, 2021; Noonan et al., 2013). More challenging semantic tasks also recruit domain-general control regions within the bilateral multiple-demand network (MDN; Duncan, 2010); however, SCN is thought to be at least partially distinct from MDN, since the peak SCN response in left anterior and ventral IFG lies outside the MDN (Gao et al., 2021; Wang et al., 2020). Given this research, we would anticipate that if creativity reflects the capacity to retrieve semantic information in a more unique way, activation within the SCN should be critical.

Retrieval from episodic memory can also be largely uncontrolled, or constrained to suit the task demands, and there are shared neurocognitive features of automatic and controlled retrieval across semantic and episodic tasks (Barredo et al., 2015; Burianova & Grady, 2007; Irish & Vatansever, 2020; Kim, 2016; Rajah & McIntosh, 2005). Default mode (DMN) regions associated with information integration (Irish & Vatansever, 2020; Lanzoni et al., 2020) and memory-guided cognitive states (Murphy et al., 2018) show common recruitment during both semantic and episodic retrieval (Burianova et al., 2010; Kim, 2016). Left angular gyrus (AG) in the core DMN may be an integrator of multimodal information across both semantic and episodic memory (Bonnici et al., 2016; Carota et al., 2021; Ramanan et al., 2017; Seghier, 2012). In addition, the left inferior frontal gyrus within the SCN shows a stronger response to both weakly associated words in semantic memory, and for words paired together fewer times in episodic memory (Vatansever et al., 2021). Damage to this site is associated with deficits in retrieving weaker semantic and episodic relations, and difficulties when these sources of information are in conflict (Stampacchia et al., 2019; 2018).

Although research has shown that creativity draws on both DMN (Beaty et al., 2015; 2016; 2014; 2020; Evans et al., 2020; Frith et al., 2020; Marron et al., 2018) and control regions (e.g., MDN), including lateral frontal cortex and dorsomedial prefrontal cortex within the SCN (Abraham et al., 2012; Chen et al., 2020; Gonen-Yaacovi et al., 2013), both semantic and episodic memory draw on these networks: this leads to uncertainty about how we generate creative patterns of thought using these long-term memory representations. The highly creative brain shows a fine balance between integration and segregation of control networks and DMN, while the less creative brain is dominated by motor and visual processing (Zhuang et al., 2021). Moreover, individuals with strong and flexible connectivity between executive networks and DMN score more highly on tests of intelligence (Sripada et al., 2019) and produce more creative ideas (Beaty et al., 2015; 2016; 2014; 2017; 2019). SCN regions are argued to be important for the interaction between executive and DMN regions, because they fall at the juxtaposition of DMN and MDN (Wang et al., 2020); SCN is unique in showing shared intrinsic and structural connectivity to both DMN and MDN which are often anti-correlated (Davey et al., 2016). Recently, research has explored functional subdivisions within DMN based on the original resting-state parcellation of 1000 brains, which provided a 17-network solution separating the DMN into three distinct subsystems (Yeo et al., 2011). A ‘dorsomedial’ DMN subnetwork – comprising nodes in dorsomedial prefrontal, lateral temporal and inferior frontal cortex – partially overlaps with the SCN (Figure 1). This observation is consistent with current accounts of DMN function that emphasise the role of this network in information integration and internally-oriented cognition across both controlled and less constrained contexts (Braga et al., 2013; Crittenden et al., 2015; Konishi et al., 2015; Krieger-Redwood et al., 2016; Lanzoni et al., 2020; Leech et al., 2011; Smallwood et al., 2021; Sormaz et al., 2018; Wang et al., 2021; Wens et al., 2019). In contrast, a ‘core’ DMN subsystem shows greater task-related deactivation, particularly during challenging decisions, and no overlap with SCN (Figure 1). These DMN subsystems have been differentially implicated in semantic (dorsomedial DMN) and episodic (core) processes (Zhang et al., 2021).

**Figure 1.**
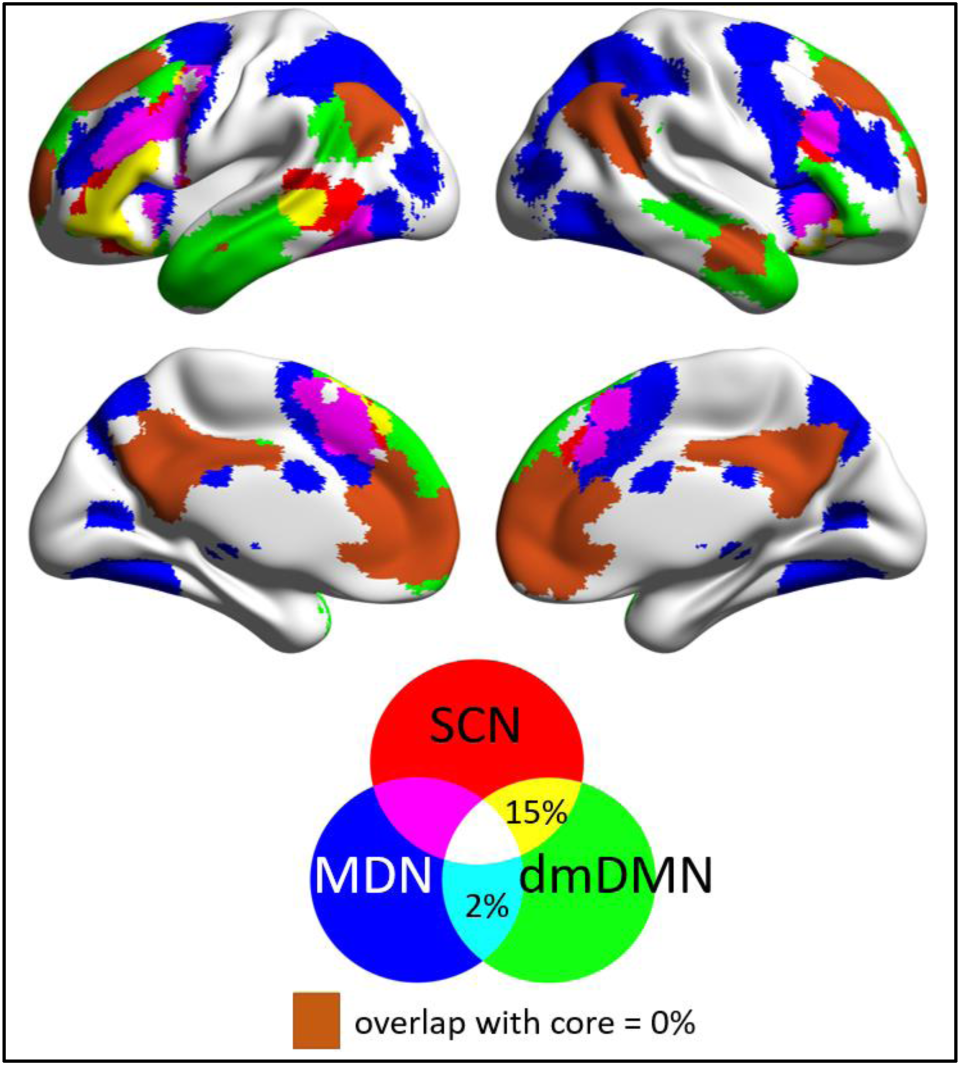
The dorsomedial DMN (dmDMN) subsystem defined from a 17-network parcellation of resting-state connectivity by Yeo et al. (2011) overlaps with the semantic control network (SCN), and minimally overlaps with the multiple demand network (MDN). In contrast, the core DMN subsystem shares no overlap with either control network.

In this study, we asked how controlled and automatic aspects of semantic and episodic retrieval support the generation of highly creative and more stereotypical verbal associations. We presented pairs of words parametrically varying in their strength of association, from strongly-related to unrelated trials. While both semantic and episodic memory representations might support the identification of links between items, the availability of relevant information in semantic and episodic memory might differ across strongly, weakly and unrelated word pairs: for example, in the absence of strong semantic links, participants might fall back on past episodes, or alternatively, in the absence of common linking episodes, participants might identify links mediated by specific semantic features. In addition, the neurocognitive processes that support controlled retrieval (from either semantic or episodic memory) might be crucial in generating creative responses, since engaging control mechanisms can promote non-dominant patterns of retrieval, while retrieval that is well-aligned across both semantic and episodic memory stores may be associated with low control demands. Accordingly, we tested five intersecting hypotheses about the neural basis of verbal creativity: (i) creative verbal behaviour will relate to specific components of long-term memory; (ii) the generation of creative links will be associated with divergence across semantic and episodic stores such that only one aspect of long-term memory drives the response; (iii) in these circumstances, control processes will be recruited to constrain retrieval from long-term memory, allowing the production of non-dominant information; (iv) this creative behaviour will be associated with recruitment of the left-lateralised SCN; and (v) the degree of creativity will modulate recruitment across distinct subsystems of DMN, since the dorsomedial subsystem, but not the core DMN, overlaps with the SCN.

## Method

### Participants

#### Task-based fMRI

We recruited 36 participants (23 females, mean age = 22 years, range = 19-32 years). None of the participants had a history of psychiatric or neurological illness, drug use that could alter cognitive functioning, severe claustrophobia, or pregnancy. All volunteers provided written informed consent and were debriefed after data collection. Ethical approval was obtained from Ethics Committees in the Department of Psychology and York Neuroimaging Centre, University of York. All participants were right-handed, native English speakers with normal/corrected vision, and were compensated for their time with payment and/or course credit. This study was not pre-registered in a time-stamped, institutional registry prior to the research being conducted.

Three participants were not included in data analysis (one withdrew during scanning, one had an anomaly in MRI, and one had missing volumes in MRI). Two further participants were removed post-analysis, one due to poor behavioural performance (no link made on 32% of trials) and one for excessive movement in two out of three runs (> 1.2mm absolute). Therefore, the final sample included 31 participants (21 females, mean age = 22 years, range = 19-32 years).

All data included in the analyses presented in this manuscript have no (absolute) movement greater than 1.2mm. For two of the 31 participants in our final sample, 2 out of 3 runs of data are included, due to excessive movement on one of the runs.

### Task Materials and Procedure

#### Pre-scan behavioural tasks

Before the neuroimaging session, participants practiced link formation at home. They were given fifteen word-pairs ranging in strength of association and were asked to generate a link between the words, type this into an answer field, and then rate the link they formed on a 1-4 scale (the same scale used in the scanner). This allowed participants to familiarise themselves with the paradigm, and provided a check that they understood the paradigm.

Participants also performed the unusual uses task (UUT). The unusual uses task (UUT) is a standard assessment of creativity, in which participants are asked to name as many uses as they can for a given object. An initial screen presented the following instructions: “In this task you will be presented with the name of an object for 10 seconds. // You will then be taken to a blank screen where you are required to list as many uses for that object as you can think of in 2 minutes. // This will be repeated for 3 different objects.” The three objects were: brick, newspaper, shoe. Participants typed their uses in the answer box, which stayed on screen for the full duration of 2 minutes, after which the next object appeared for 10 seconds, followed by the 2-minute generation screen. Both of these paradigms were presented using Qualtrics (www.qualtrics.com).

#### fMRI behavioural tasks

Each trial lasted 13.5 seconds and was structured as follows. Participants were presented with word pairs on the screen for 4.5 seconds (Figure 2**Error! Reference source not found**.). During this period, participants were tasked with identifying a link between the words. They were given no specific instructions to be creative. The words varied in the degree to which they had a pre-existing semantic link. We manipulated semantic association strength between the word pairs using word2vec (Mikolov et al., 2013) to identify trials ranging from completely unrelated (i.e., minimum word2vec = -.05) to strongly related (i.e., maximum word2vec = .72). Word2vec uses word co-occurrence patterns in a large language corpus to derive semantic features for items, which can then be compared to determine the degree of their relationship. Following link generation for each trial, there was a fixation period of .5 seconds, followed by 1 s to rate the strength of the link that was identified. Participants were specifically instructed to rate the strength of the link they had made, and not the pre-existing strength of association between the words. They provided ratings on a 1-4 scale (weak – strong), using their right hand, or alternatively pressed a button with their left hand to indicate no link was made. Following the link rating, participants performed a series of left-right chevron decisions (details below). There were 144 word-pair trials in total, presented across three runs, and these were pseudo-randomly assigned such that each run contained an even number of high, medium, low and unrelated associate pairs.

**Figure 2.**
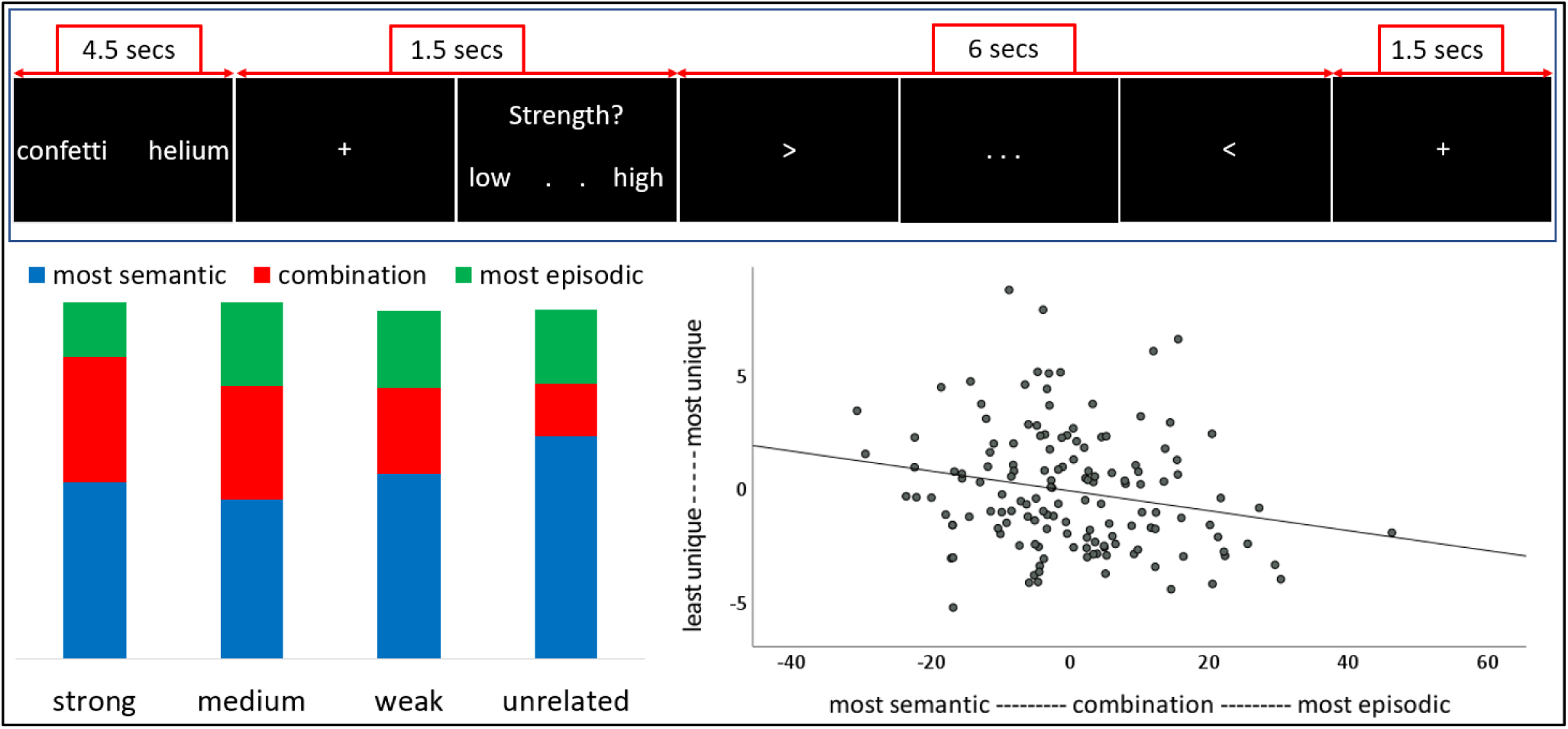
Top: Task schematic. Participants covertly generated a link between the two words; next they rated the strength of the link that they had formed, and then engaged in a series of fast-paced left-right chevron decisions. Bottom Left: The distribution of memory types used to generate associations for strong, medium, weak and unrelated trials. Participants rated the memory used on a 5-point scale, with most semantic on one end and most episodic on the other. Combination refers to trials drawing on both episodic and semantic memory. As the semantic relationship between the words became more distant, participants relied more on semantic memory, but when the pre-existing relationship between words was stronger, participants could make use of both semantic and episodic memory. Bottom Right: Partial plot (controlling for word2vec) of the relationship between the uniqueness of the response and rated engagement of semantic-to-episodic memory. The values in this plot are based on an item analysis of each word pair (the y-axis shows the number of response types given for each word pair; the x-axis is the shows the engagement of semantic to episodic memory).

#### Post-scan behavioural tasks

Immediately following the scanning session, post-scan behavioural testing included a series of questions about in-scanner performance. Participants were asked to: i) recall and describe the link that they formed in the scanner; ii) rate the strength of the link that they formed on a 5 point scale (for the second time, since the same judgement was made in the scanner); iii) report the degree to which the link on that trial relied on semantic to episodic memory (a 5 point scale with semantic on one end, combination in the middle and episodic at the other end), and iv) rate their confidence in their recall of the association they formed in the scanner (5 point scale). There was a high degree of overlap between: participants’ in-scanner ratings of the link they made and word2vec (Pearson *r* = .83, p < .001); word2vec and post-scan ratings (Pearson *r* = .82, p < .001); and in-scanner and post-scan ratings (Pearson *r* = .98, p < .001).

After these questions, participants also performed a standard three alternative-forced-choice semantic association task (e.g., carrot - <u>dinner</u>, celebrity, television). This consisted of 120 trials (60 strong associations; 60 weak associations), presented in four mini-blocks. The task started with an instruction screen, which had no deadline and participants could initiate the start of the task at their own pace. Each trial remained on screen until an answer was given (maximum duration 3 seconds), after which the next item was presented. The target and distractors were presented first, and then after 900ms the probe appeared and participants could make their response. Responses were not recorded for 4 participants; therefore, this task was not included in any data analysis.

### fMRI Task Procedure

Before entering the scanner, participants re-practiced the link formation task for two trials, stating their retrieved associations aloud, with feedback from the experimenter. They then completed 25 practice trials on a computer using the same presentation format as the task in the scanner, with 4.5 seconds for each word-pair followed by the link rating question, and chevron trials.

The MRI session included a localiser scan, 3 functional runs (11 minutes and 45 seconds each), and a structural T1 scan following completion of the 3 functional runs. We used a slow event-related design, with 7.5 seconds between trials; 6 seconds were filled with the chevron task – participants indicated whether the chevron faced left or right, with 10 chevrons presented across this 6 second block). There was then 1.5 seconds of fixation to alert the participant to the upcoming trial (Figure 2). Halfway through each run participants had 30 seconds of rest to help maintain focus.

### FMRI Acquisition

Whole brain structural and functional MRI data were acquired using a 3T Siemens MRI scanner utilising a 64-channel head coil, tuned to 123 MHz at York Neuroimaging Centre, University of York. A Localiser scan and 3 whole brain functional runs were acquired using a multi-band multi-echo (MBME) EPI sequence, each 11.45 minutes long (TR=1.5 s; TEs = 12, 24.83, 37.66 ms; 48 interleaved slices per volume with slice thickness of 3 mm (no slice gap); FoV = 24 cm (resolution matrix = 3×3×3; 80×80); 75° flip angle; 455 volumes per run; 7/8 partial Fourier encoding and GRAPPA (acceleration factor = 3, 36 ref. lines); multi-band acceleration factor = 2). Structural T1-weighted images were acquired using an MPRAGE sequence (TR = 2.3 s, TE = 2.26 s; voxel size = 1×1×1 isotropic; 176 slices; flip angle = 8°; FoV= 256 mm; interleaved slice ordering).

### Multi-echo data pre-processing

This study used a multiband multi-echo (MBME) scanning sequence to optimise signal from medial temporal regions (e.g., ATL, MTL) while also maintaining optimal signal across the whole brain. We used TEDANA (version 0.0.7) to combine the images (https://tedana.readthedocs.io/en/latest/outputs.html; Kundu et al., 2013; Posse et al., 1999). Before images were combined, some pre-processing was performed. FSL_anat (https://fsl.fmrib.ox.ac.uk/fsl/fslwiki/fsl_anat) was used to process the anatomical images, including re-orientation to standard (MNI) space (fslreorient2std), automatic cropping (robustfov), bias-field correction (RF/B1 – inhomogeneity-correction, using FAST), linear and non-linear registration to standard-space (using FLIRT and FNIRT), brain extraction (using FNIRT, BET), tissue-type segmentation (using FAST) and subcortical structure segmentation (FAST). The multi-echo data were pre-processed using AFNI (https://afni.nimh.nih.gov/), including de-spiking (3dDespike), slice timing correction (3dTshift; heptic interpolation), and motion correction (3dvolreg applied to echo 1 to realign all images to the first volume; these transformation parameters were then applied to echoes 2 and 3; cubic interpolation). The script used to implement the pre-processing TEDANA pipeline is available at OSF (https://osf.io/ydmt4).

### fMRI data analysis

First, second and group-level analyses were conducted using FSL-FEAT version 6 (FMRIB’s Software Library, www.fmrib.ox.ac.uk/fsl; Jenkinson et al., 2002; Smith et al., 2004; Woolrich et al., 2009). The TEDANA outputs (denoised optimally combined timeseries) registered to the participants’ native space were submitted to FSL, and pre-processing included high-pass temporal filtering (Gaussian-weighted least-squares straight line fitting, with sigma = 50s), linear co-registration to the corresponding T1-weighted image followed by linear co-registration to MNI152 standard space (Jenkinson & Smith, 2001), spatial smoothing using a Gaussian kernel with full-width-half-maximum (FWHM) of 6 mm, and grand-mean intensity normalisation of the entire 4D dataset by a single multiplicative factor.

Pre-processed time series data were modelled using a general linear model correcting for local autocorrelation (Woolrich et al., 2001). We used an event-related parametric design – the linear model included three experimental conditions as parametric variables (start time, duration, and a mean-centred parametric regressor for each trial). Our analysis focussed on the effect of (1) word2vec (i.e., a measure of semantic control based on the degree of pre-existing semantic relatedness between the two words); (2) uniqueness (i.e., the degree to which each participant’s response was unique/creative (see behavioural results); and (3) the degree to which the participant used episodic memory to form the link. Other EVs were: (4) mean activation for the trial (start, duration and weighting of 1), (5) participant judgment of link made (1.5secs), (6) rest (30 seconds of rest, occurring mid-way through each run), (7) fixation and (8) the first 2 chevron trials. We modelled the first 2 trials of the implicit baseline (chevron task) in order to account for the switch cost (i.e., moving from link formation to fast-paced chevrons). We did not include any motion parameters in the model as the data submitted to these first level analyses had already been denoised as part of the TEDANA pipeline (Kundu et al., 2012). All group-level analyses were cluster corrected using a *z*-statistic threshold of 3.1 to define contiguous clusters (Worsley, 2001) and then cluster corrected for multiple comparisons at *p* < 0.05 FWE; the group-analyses were run within a liberal grey-matter mask (40% probability of GM). Conjunction analyses were run using FSL’s ‘easythresh_conj’ tool across all of the task conditions (weak association, strong association, episodic, unique). All maps generated are freely available at Neurovault (https://identifiers.org/neurovault.collection:8799).

## Results

### Behavioural Results

We first examined participants’ ratings of whether they used semantic memory, episodic memory or a combination to retrieve each link (see Figure 2). Participants reported that they primarily identified links using semantic memory (Greenhouse-Geisser corrected, F(1.5, 45) = 24.8, p < .001; Figure 2). To understand how the balance of semantic to episodic retrieval varied with the relatedness of the words, we split the trials into four categories based on word2vec: Strong, Medium, Weak, Unrelated. The type of memory used to establish a link varied significantly across these categories (memory type by association strength interaction, Greenhouse-Geisser corrected, F(1.8, 55) = 5.9, p = .006; Figure 2). For distant items, participants were more reliant on semantic memory (Greenhouse-Geisser corrected, F(1.2, 36) = 4.03, p = .046; Figure 2) and were less likely to use a combination of semantic and episodic memory (Greenhouse-Geisser corrected, F(1.4, 41) = 14.88, p < .001; Figure 2). Episodic memory only contributes to our identification of meaningful connections between words when the items are strongly related and we are likely to have encountered them together. In these circumstances, when we generate links for highly related words, semantic and episodic information is likely to coherently support link generation, since these items are likely to have been encountered together during an everyday event which is then available in episodic memory, as well as having related meanings.

Using participants’ post-scan recall of the links that they formed, we analysed the uniqueness of each response. These values were expressed as a proportion of the total sample who gave that particular response, ranging from .03 (a minimum of 1/31 participants) to 1 (a maximum of 31/31 participants). Word pairs with the lowest word2vec scores (i.e., unrelated) had the greatest variety of responses across participants, while word-pairs with strong association strength tended to generate similar responses in different individuals (*Pearson r* = -.61, *p* < .001; Figure S5). Furthermore, we found that as participants relied more on episodic memory, their responses became less unique (partial correlation controlling for word2vec; *Pearson r* = -.21, *p* = .013; Figure 2, right-hand side). Additional behavioural analyses pertaining to participant confidence in recall, can be found in the supplementary materials (76% of recall was self-rated as highly confident (3 or 4 on a 0-4 scale), while only 12% of recall was rated as low confidence (0 or 1 on 0-4 scale); SM section: Behavioural Data).

### fMRI results

There were clear differences in the neurocognitive processes that underpinned the retrieval of links between words, depending on their associative strength. Strongly related word pairs (with high word2vec values) were often linked in a stereotypical way, which was common across participants. The parametric effect of strongly linked items was associated with greater activation in swathes of medial parietal and medial occipital cortex (Figure 3). Peak responses were observed in right inferior lateral occipital cortex, left postcentral gyrus extending into precentral and superior parietal lobule, right and left parietal operculum and right and left central operculum. These effects overlapped with visual, motor and ventral attention networks (Figure S4; Yeo et al., 2011). Decoding this map using Neurosynth identified terms consistent with a role in less constrained, more automatic cognition, such as ‘sensorimotor’, and ‘resting’ (Figure S3). In contrast, as participants retrieved links between words that were more distantly related, activation increased in semantic control and multiple-demand networks, with a stronger response in left inferior frontal gyrus, dorsomedial prefrontal cortex, bilateral insula, and left posterior middle and inferior temporal gyri (Figure 3; Figure 4). There were also increases in activation beyond control networks, in posterior fusiform gyrus, right occipital pole and cerebellum. Functions associated with this map, decoded using Neurosynth, encompass executive terms such as ‘working memory’ and ‘demands’ and language terms such as ‘semantic’, ‘language’ and ‘reading’ (Figure S3).

**Figure 3.**
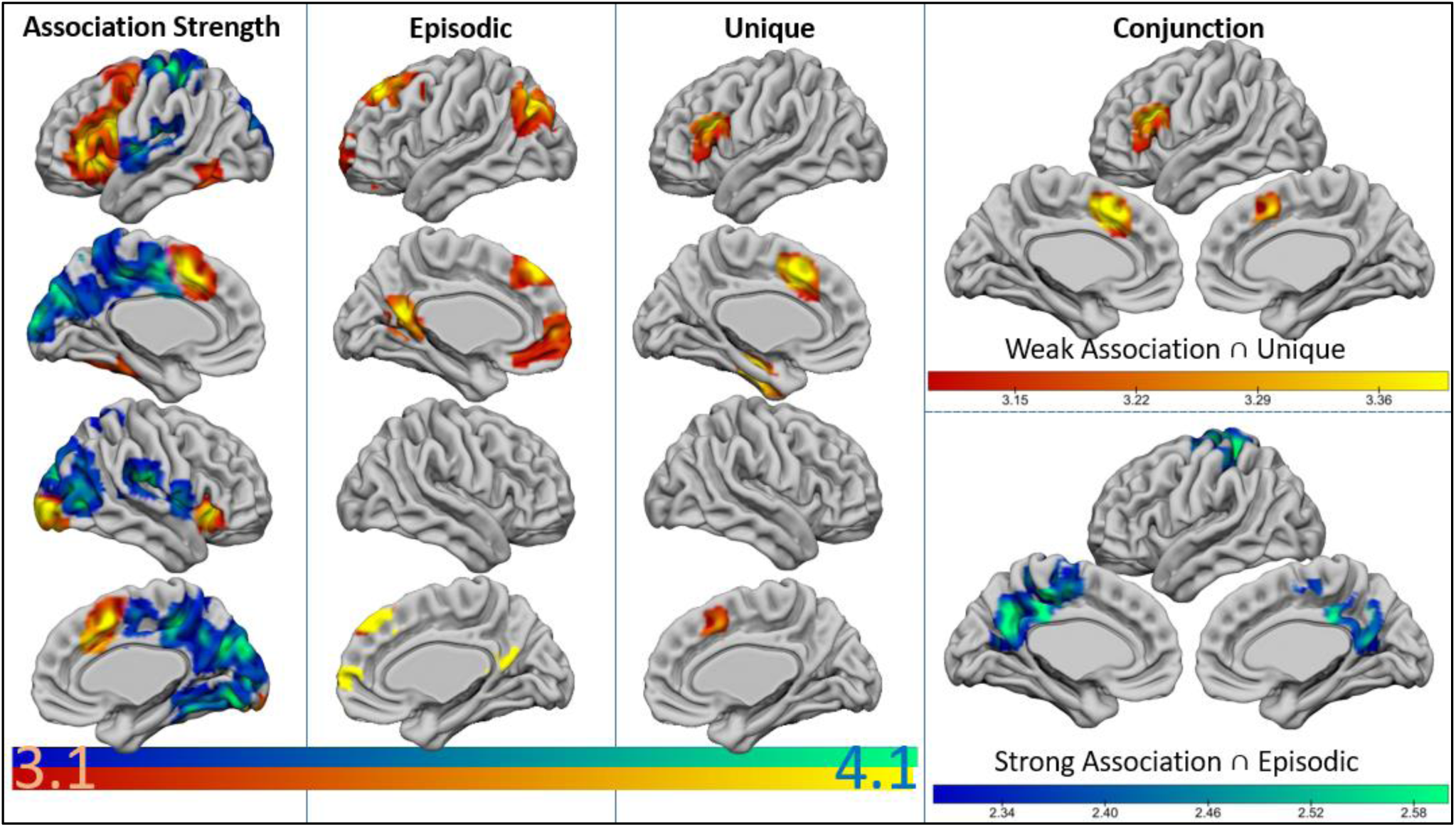
fMRI activation for the parametric effects (z ≥ 3.1, p ≤ .05). The first column shows thresholded activation for weak (red-yellow) to strong (blue-green) association strength as measured by word2vec ratings. The second column shows areas in which activation increased as participants became more reliant on episodic memory. The third column displays activation associated with more unique responses. The fourth column shows the activation across pairs of regressors, with conjunctions observed for weak associations and unique responses, as well as for strong associations and more episodic retrieval. There were no conjunctions when these conditions were recombined (i.e. no conjunction of weak associations and episodic retrieval, or strong associations and unique responses). Even a more lenient analysis (z > 2.3, p < .05) designed to minimise Type II errors confirmed this pattern of selective conjunctions between (i) weak association ∩ uniqueness and (ii) strong association ∩ episodic (and not the reverse).

**Figure 4.**
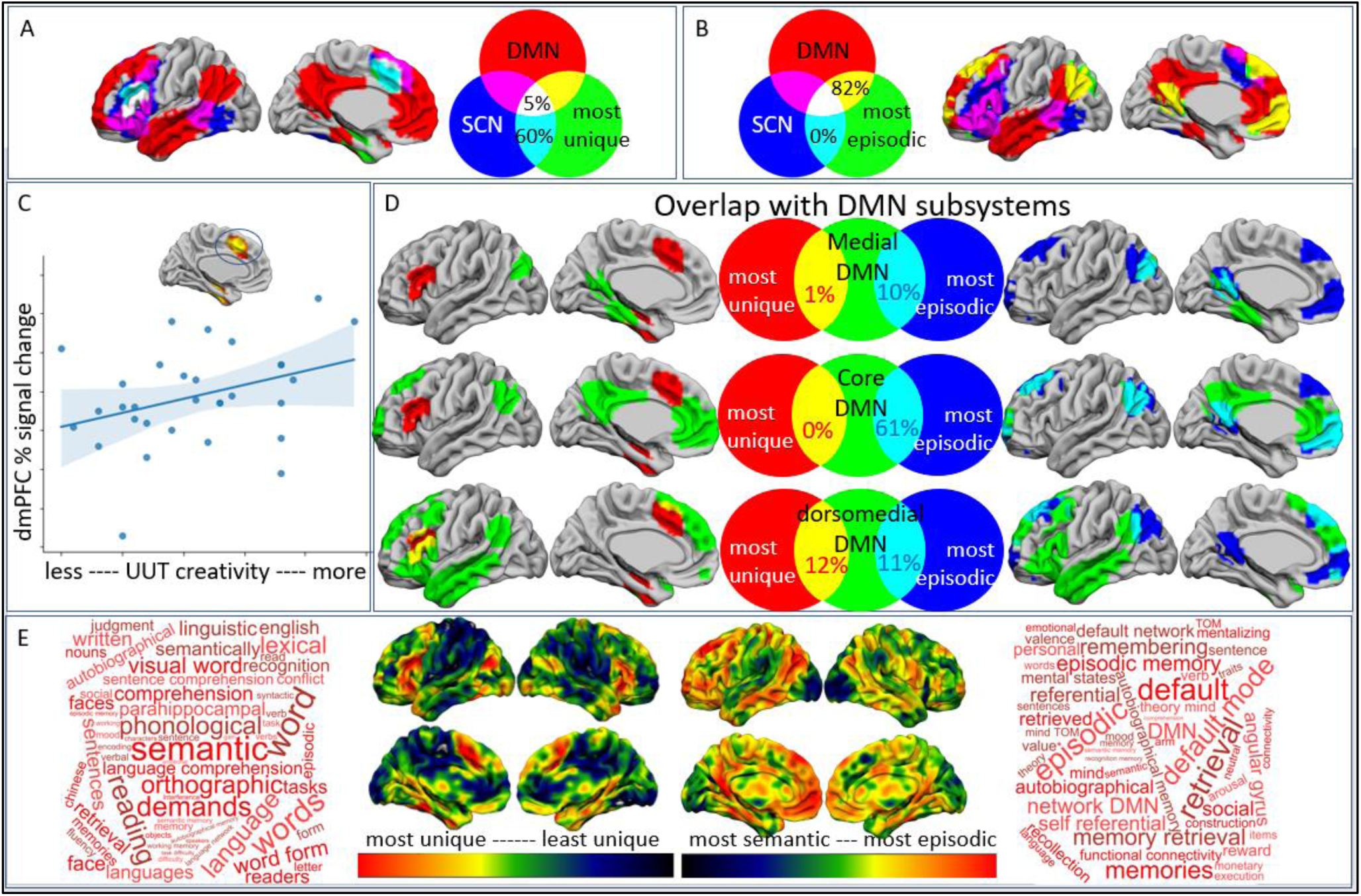
Top row: Overlap of activation for parametric effects for most unique and most episodic responses with the following two networks: semantic control (SCN from a meta-analysis of semantic control; Jackson 2020) and Default Mode (DMN from the 7-network parcellation of resting state data; Yeo et al. 2011). The Venn diagrams show the percentage of voxels for each effect that overlapped with these established networks, both unique (A) and episodic (B) responses fall within non-overlapping parts of the DMN. Middle panel: (C) The relationship between increased activitation for more unique responses in dmPFC and more creative performance on the Unusual Uses Task (UUT) outside of the scanner. (D) Overlap of activation for parametric effects, with the Default Mode Network (DMN) subsystems: medial (Yeo 15), core (Yeo 16) and dorsomedial (Yeo17). The Venn diagrams show the percentage of overlapping voxels for each effect with these established networks. Bottom panel (E): Unthresholded activation maps showing the continuous response associated with the parametric regressors. The word-clouds are derived from a Neurosynth meta-analysis of these maps.

Our behavioural analysis showed that trials of different associative strengths elicit responses that vary in their degree of uniqueness across individuals. Weakly-associated word pairs tend to elicit more diverse associations across participants, suggesting this pattern of retrieval places higher demands on processes that support creativity. A regressor examining changes in activation as participants’ responses became more unique revealed greater activation in left inferior frontal gyrus and dorsomedial prefrontal cortex (dmPFC). Both of these clusters overlapped with areas of the semantic control network that showed more activation for weak associations (Figure 3; Figure 4A). A formal conjunction analysis confirmed this pattern of overlap (Figure 3). Cognitive decoding using Neurosynth revealed an overlap with both semantic and executive responses – identifying terms such as ‘semantic’, ‘demands’, etc. (Figure 4E). Since greater activation within the semantic control network for more unique responses was identified in a model that also included word2vec as a regressor, this analysis suggests that activation within the semantic control network can be observed in response to more creative responses. There was also greater activation in temporal fusiform cortex for more unique responses, which did not overlap with the effect of presenting weaker associations (reported above). This finding additionally suggests that anterior parts of the medial temporal lobe support the ability to generate a novel connection between two words.

A final regressor examined how the neural response during link generation varied as a function of reliance on episodic memory. On trials in which participants indicated that they were drawing more on episodic memory, stronger left-lateralised activation was seen in angular gyrus, ventral and dorsal clusters within anterior cingulate cortex extending into frontal pole and superior frontal gyrus, and in posterior cingulate cortex extending into retrosplenial cortex (Figure 3). These regions were largely overlapping with the DMN (Figure 4B) and included sites implicated in episodic memory. Cognitive decoding of the map in Neurosynth elicited terms such as ‘autobiographical’, ‘retrieval’ and ‘episodic’ (Figure 4E). The uniqueness of the response (see above) did not overlap with this effect of reliance on episodic memory: more creative links, generated by only a few participants, elicited activation in anterior aspects of the medial temporal lobes, while the episodic memory regressor was associated with greater posterior medial temporal and parietal activation. However, the reliance on episodic memory did overlap with the effects of strong associative strength (captured by word2vec), particularly in medial parietal regions. A formal conjunction analysis confirmed this pattern of overlap (Figure 3).

We identified DMN activation when participants generated both more episodic and more unique responses, yet there was an absence of overlap between these two regressors. We therefore assessed the contribution of three previously-described DMN subsystems (medial, core, dorsomedial; Yeo et al. 2011) to link formation. Greater availability of episodic information during the generation of verbal associations primarily activated the core DMN, while also eliciting activation in the dorsomedial and medial DMN subsystems (Figure 4D). In contrast, when links were formed for weakly associated words, and when responses were more unique, DMN activation fell within the dorsomedial DMN, with little or no activation in the other subsystems (Figure 4D; Figure S3). In order to assess whether activation within these DMN subsystems was significantly different across our regressors, we extracted the percentage of each participant’s activation map (thresholded at *z* = 2.3) that fell within each DMN subsystem (dorsomedial, medial, core) for each regressor (word2vec, uniqueness, episodic). Episodically-mediated trials overlapped with significantly more of the core DMN than unique (t(30) = -2.481, *p* = .019) and weak-associate (t(30) = 3.551, *p* = .001) responses. The percentage of voxels within each map falling within the dorsomedial DMN subsystem did not significantly differ across episodic, weakly associated, or unique link generation regressors (F(2, 60) < 1). There was also no difference between these regressors within the medial subsystem (F(2, 60) = 2.1, p = .13).

#### Correlation with Unusual Uses Task

Our analyses revealed activation in left inferior frontal gyrus, dorsomedial prefrontal cortex and temporal fusiform cortex for more unique word-pair link formation. In order to assess whether the activation during link formation predicted individual differences in creativity on a more standard assessment, we determined whether the strength of the uniqueness effects in these regions was associated with performance on the Unusual Uses Task. In a regression analysis predicting Unusual Uses performance from the activation in each of these clusters simultaneously, we found that increased activation in dmPFC during the generation of unique links correlated with better performance on Unusual Uses Task (F(1, 27) = 6.95, *p* = .014; Figure 4C). Activation within the other clusters did not make a unique contribution to creativity (inferior frontal gyrus: F(1,27) = 2.545, *p* = .12; temporal fusiform: F(1, 27) = 2.2, *p* = .15).

As a control, we also confirmed that activation in a cluster adjacent to dmPFC (elicited by the episodic regressor) did not predict Unusual Uses performance (F(1, 26) = 1.3, p = .3; see supplementary materials: Correlations with Unusual Uses Task Performance). These analyses establish the link between activation within the semantic control network and creativity, and also suggest that this involvement of controlled semantic retrieval in creativity is independent of the recruitment of episodic memory to support link generation.

## Discussion

This study investigated the contribution of brain networks supporting semantic and episodic retrieval as participants generated links between words that varied in their degree of association along a continuum from strongly related (lowest creativity) to unrelated (highest creativity). In this way, we were able to test hypotheses about the contribution of different long-term memory processes and neural networks related to creative and automatic verbal behaviour. Participants largely relied on semantic information to identify links between words, although episodic memory contributed to link generation for strongly-associated trials. Creative connections were generated through flexible and controlled retrieval of less dominant semantic information – with greater recruitment of the semantic control network (SCN) in left inferior frontal gyrus and dorsomedial prefrontal cortex (dmPFC) when unusual links were generated. The functional network supporting verbal creativity partially overlapped with the dorsomedial DMN subsystem and unique responses generated activation in this subsystem. In contrast, strong associations aligned across both episodic and semantic aspects of long-term memory and supported by the relatively automatic patterns of retrieval were associated with activation in core DMN. These results support the notion that creativity emerges from an interaction of memory and control processes (Zhuang et al., 2021) – but go beyond the state of the art to reveal the specific neurocognitive processes that drive activation in control and DMN networks.

When two items are strongly associated, people are more likely to have episodic memories of their interaction or relationship – and therefore they can generate links relying on both episodic and semantic memory. This fits with the emerging literature demonstrating that these two memory systems draw on distinct yet interacting long-term stores, and share common automatic and controlled retrieval pathways (Burianova & Grady, 2007; Burianova et al., 2010; Irish & Vatansever, 2020; Rajah & McIntosh, 2005; Vatansever et al., 2021). The activation associated with the use of episodic memory to generate links between items in the current study overlapped with regions previously implicated in more automatic (not controlled) aspects of episodic retrieval, including in left AG, ventromedial prefrontal cortex, posterior cingulate/precuneus and middle and superior frontal gyri, largely within core DMN. The AG, and other regions of DMN, are purported to play a role in binding and integrating information, in both episodic and semantic memory (Bonnici et al., 2016; Ramanan et al., 2017; Seghier, 2012) and the activation seen for our task may have reflected the integration of semantic and episodic contributions to link generation. Moreover, this situation involving information integration may promote a pattern of ‘ecphory’ – i.e. strong automatic retrieval driven by highly-constrained circumstances (Renoult & Rugg, 2020).

It is unlikely that items with weak association will have been encountered together in a past event, leaving only semantic features available to form a link. While it might be assumed that semantic knowledge is broadly shared across participants (despite individual differences that reflect interests and expertise), we observed considerable variability in the links that participants formed between words, especially with greater semantic distance. The accompanying neural activation was spread across control networks and dorsomedial DMN - this activation profile was also seen when responses were more unique. The SCN sits at the intersection of dorsomedial DMN and MDN regions in the left hemisphere (Wang et al., 2021), with both structural and functional connectivity to regions within both networks (Davey et al., 2016), and is therefore well positioned to leverage these networks in support of link formation for distantly related concepts. Activation in two key SCN nodes, left inferior frontal cortex and dmPFC, was common to weak associate word-pairs and when the link generated was unique. We also saw activation in other parts of the semantic control network for weakly-associated concepts, including in inferior pMTG/ITG – a site commonly recruited by more difficult semantic judgments (Davey et al., 2015; Davey et al., 2016; Whitney et al., 2011; Whitney et al., 2012).

There was also a functional dissociation within the DMN. The core DMN subsystem, encompassing posterior and anterior cingulate cortex plus angular gyrus, showed more activation when participants relied on episodic memory, primarily when the word pairs had a strong pre-existing semantic relationship. In contrast, the dorsomedial DMN subsystem, which encompasses anterior ventral parts of inferior frontal gyrus as well as temporal and parietal regions, responded during the retrieval of weaker and unique associations that were more reliant on semantic memory alone, with no activation in the core DMN. Large-scale meta-analyses have implicated dorsomedial DMN in conceptual processing (Andrews-Hanna et al., 2010; Andrews-Hanna et al., 2014), while the core and medial DMN subsystems show greater recruitment during episodic memory (Huijbers et al., 2011; Sestieri et al., 2011), past and future autobiographical thought and self-referential processing (Andrews-Hanna et al., 2010; Andrews-Hanna et al., 2014; Chiou et al., 2020; Zhang et al., 2021). In addition, the dorsomedial DMN but not the core DMN (as defined by a parcellation of resting-state fMRI of 1000 brains) overlaps with the functionally-defined SCN; this provides further evidence that dorsomedial DMN supports both automatic and controlled aspects of semantic cognition.

The activation pattern across DMN and control regions is consistent with resting-state and task-based functional studies of creativity: these networks play a complementary role in the generation and evaluation of ideas (Beaty et al., 2016; Beaty et al., 2019; Frith et al., 2020; Xie et al., 2021; Zhuang et al., 2021). The increased activation in key nodes of the SCN during unique responses might reflect the way that creativity emerges from core cognitive processes involving memory, attention and executive control (Abraham & Bubic, 2015; Abraham et al., 2012; Benedek & Fink, 2019; Frith et al., 2020; Zhuang et al., 2021). For example, Zhuang et al. (2021) suggest that coupling of DMN and executive networks is critical for creativity. The SCN is ideally situated to support this network interaction, as it is physically located between aspects of DMN and MDN on the cortical surface. This account can therefore explain why key SCN regions, in left inferior frontal cortex and dmPFC, showed greater activation during the generation of unique responses. Leveraging the SCN may allow activation to be directed towards unusual and non-dominant features and associations of concepts, so that a unique link can be identified. Furthermore, the observation that dmPFC activation was linked to better performance on the Unusual Uses Task outside the scanner, corroborates previous studies demonstrating dmPFC as a key player in creative cognition (Boccia et al., 2015; Gonen-Yaacovi et al., 2013).

A limitation of this study is that we cannot model the emergence of creative idea generation in a dynamic way: we used a slow-event-related design to maximally separate activation across trials, and are unable to model activation at different time-points in the generative process – for example, to investigate whether aspects of DMN couple with ventral attention versus control networks during initial retrieval and later elaboration (cf. Beaty et al., 2015). Our results revealed activation in executive and default mode networks, but little activation in the ventral attention network (VAN), for both weak associations and more unique links (only strong association word pairs elicited VAN activation, consistent with the detection of salient associations between items in long-term memory that were sufficient for performing the task; Figure S2). Secondly, while two of the clusters associated with more unique responses fell within the SCN (left inferior frontal gyrus and dorsomedial prefrontal cortex), one did not, in temporal fusiform cortex. However, this site is often associated with semantic processing (Chrysikou & Thompson-Schill, 2011; Ding et al., 2016; Ellamil et al., 2012; Mion et al., 2010; Shen et al., 2017). Fusiform gyrus, alongside left inferior frontal cortex, is reported to show maximal activity for unrelated word pairs, and least activity when identical words are repeated (Wheatley et al., 2005), similar to our study, where more disparately related concepts elicited greater fusiform activation (posterior for weak associations, and anterior for more unique generation). Fusiform cortex has also been implicated in the formation of new associations, when participants are required to generate uncommon uses for objects (Chrysikou & Thompson-Schill, 2011). Shen et al. (2017) propose that the fusiform gyrus has at least two roles in creative problem solving: (i) ‘gestalt-like’ processing of feature conjunctions and (ii) perspective taking (i.e., taking a different/new perspective other than the most salient meaning of a word, for example, by thinking of a shoe as a flower pot rather than an item of clothing).

Finally, while previous studies have increased creativity using episodic induction prior to creative idea generation (Beaty et al., 2020; Madore et al., 2015; 2016a; 2016b; 2019), episodic memory was linked to *less* unique responses in our study. This suggests that the neurocognitive basis of creative idea generation may vary with the task: here, participants were required to generate a link between two words, which required semantic processing on every trial (for example, to access the meanings of the individual words). Consequently, in our paradigm, the retrieval of episodic information on strongly-associated trials had a highly constraining effect on patterns of retrieval (since episodic and semantic sources of information were likely to be coherent on these trials). Future studies could continue to unpick the psychological processes that contribute to different types of creative behaviour, for example, by examining which aspects of semantic control (e.g., flexible retrieval, selection from amongst competing alternatives, conceptual combination, etc) correspond with convergent and divergent creativity.

In conclusion, this study asked participants to produce links between two words to establish the contribution of semantic and episodic memory to our capacity to creatively link ideas, and also examined the neurocognitive processes that underpin more unique compared with more stereotypical responses. We found that participants engaged semantic memory for more creative generation, with accompanying recruitment of the SCN. When semantic and episodic memory stores were well aligned (i.e., for more stereotypical links), activation was dominated by the DMN. Furthermore, we uncovered a dissociation within DMN during link generation. The core DMN was recruited when information from episodic and semantic memory systems was likely to be coherent, supporting information integration. In contrast, the core DMN was not implicated in the semantic control processes required for more unique ideas, but, areas within the dorsomedial DMN were; these trials were more reliant on semantic information to generate a link and were less constrained by experiences in episodic memory.

## Acknowledgements

The study was funded by the European Research Council [FLEXSEM-771863]. JS was supported by the European Research Council [WANDERINGMINDS-646927]. AH was supported by the Rosetrees Trust (A1699) and a Career Development Award from the Medical Research Council (MR/V031481/1). We would like to thank Dominika Varga, Megan Evans and Johannes Wibroe for their help with data collection.

