## SupplementaryInformation for "Creativity in Verbal Associations is Linked to Semantic Control"

### Supplementary Materials

#### Conjunction Analyses

Conjunction analyses were run using FSL's 'easythresh_conj' tool and thresholded at z = 2.3, $p < .05$. Conjunction analyses were run across all of the task conditions (low w2v, high w2v, episodic, unique). All significant conjunction results are shown in the tables below.

*Table S1 Word2Vec Low ∩ Unique*

| Cluster Index | Voxels | Label | *p* | *Z* | x | y | z |
|---|---|---|---|---|---|---|---|
| 2 | 1662 | LIFG | 0.0002 | 4.26 | -38 | 4 | 30 |
| 1 | 975 | dmPFC | 0.0085 | 4.69 | -4 | 14 | 46 |

*Table S2 Word2Vec High ∩ Episodic*

| Cluster Index | Voxels | Label | *p* | *Z* | x | y | z |
|---|---|---|---|---|---|---|---|
| 1 | 1611 | L SMA | 0.0027 | 3.52 | -22 | -22 | 62 |

[No significant conjunction between: w2v low and episodic maps; episodic and uniqueness maps; w2v high and uniqueness maps.]

A more stringent conjunction analysis thresholded at z = 3.1, $p < .05$ revealed a significant conjunction of low w2v and uniqueness, but none of the other contrasts.

*Table S3 Word2Vec Low ∩ Unique*

| Cluster Index | Voxels | Label | *p* | *Z* | x | y | z |
|---|---|---|---|---|---|---|---|
| 2 | 302 | dmPFC | 0.0068 | 4.69 | -4 | 14 | 46 |
| 1 | 257 | LIFG | 0.0144 | 4.19 | -46 | 20 | 22 |

#### Uniqueness/Episodic by Semantic Association

We split the data in half, according to semantic relationship (i.e., word2vec), to ask whether activation changed for uniqueness and/or episodic trials dependent on pre-existing semantic relationships. The figure below (Figure S1) confirms that the semantic control network is activated for more unique responses to weakly associated word-pairs, while strongly associated concepts that were linked episodically recruit core and medial DMN.

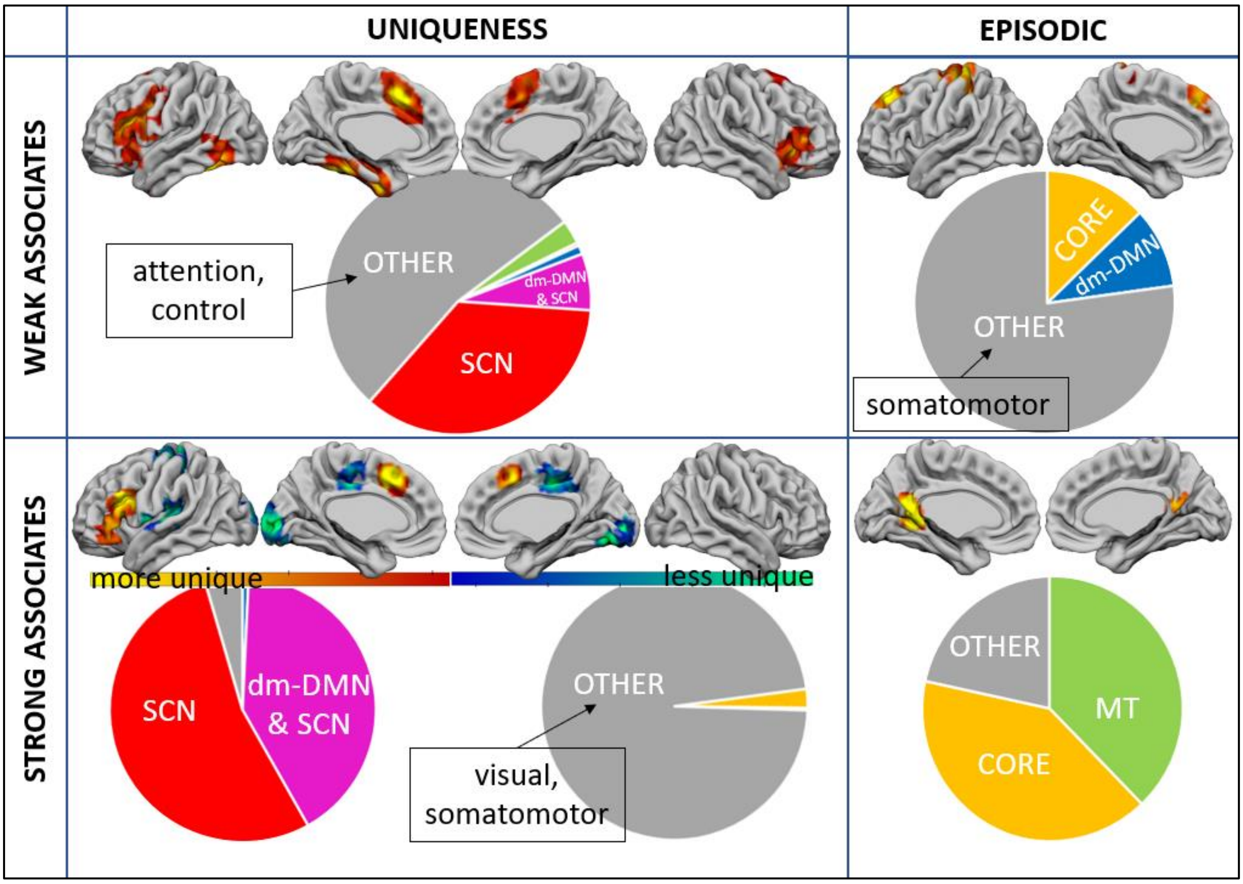


*Figure S1: Top row shows significant effects for most unique (left) and episodic (right) responses for weak associate word pairs. Bottom row shows significant effects for both most (left) and least (right) unique responses and for most episodic responses (right) for strong associate word pairs.*

Activation Overlap with Yeo Networks (17-Network Solution)

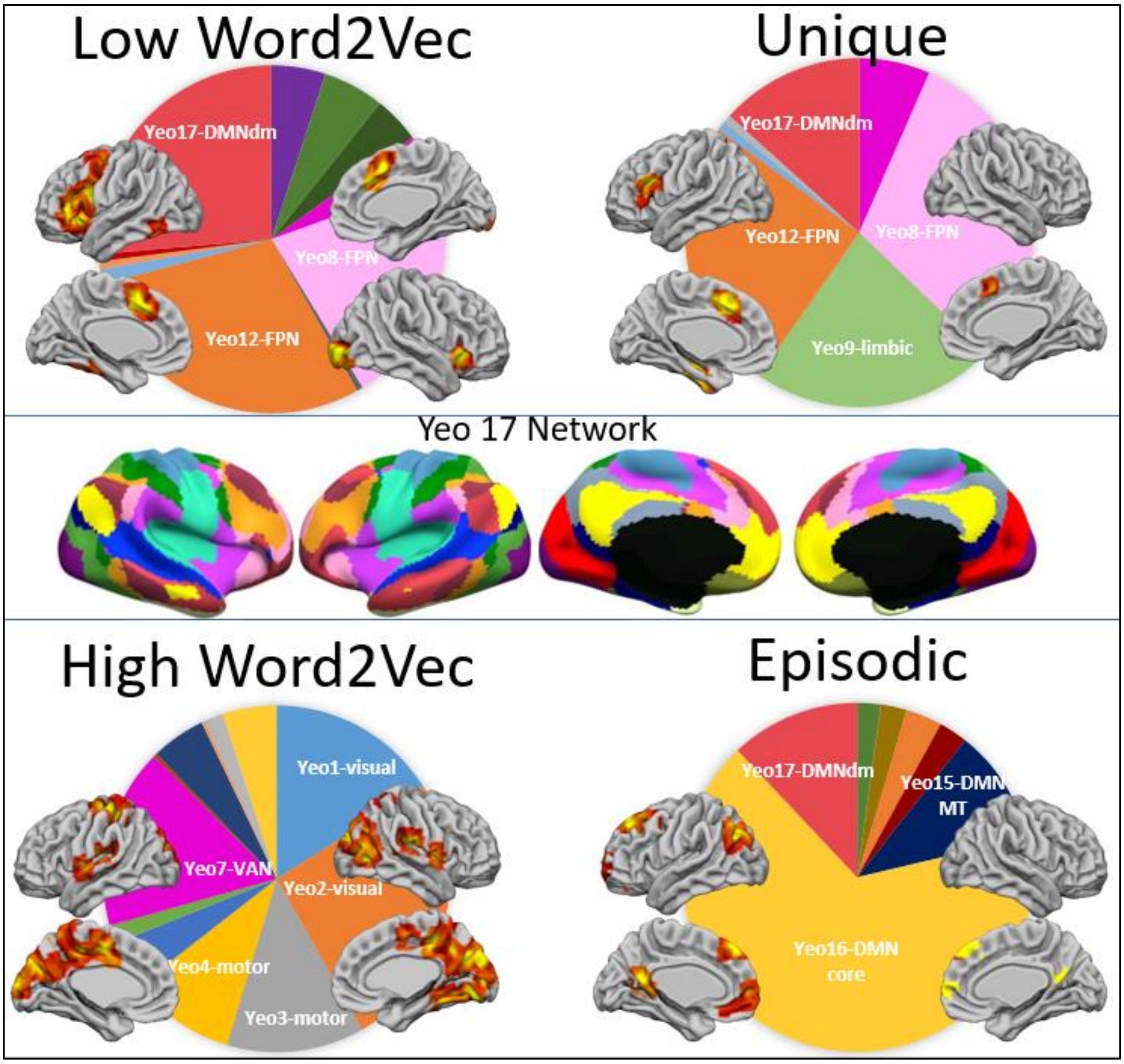


*Figure S2: Yeo Networks separated by task regressor. The brains in the middle are taken from the Yeo 17 network parcellation, colours in the pie charts relate to the colours on these brains. Brains overlaid on pie-charts show the activation for that task regressor.*

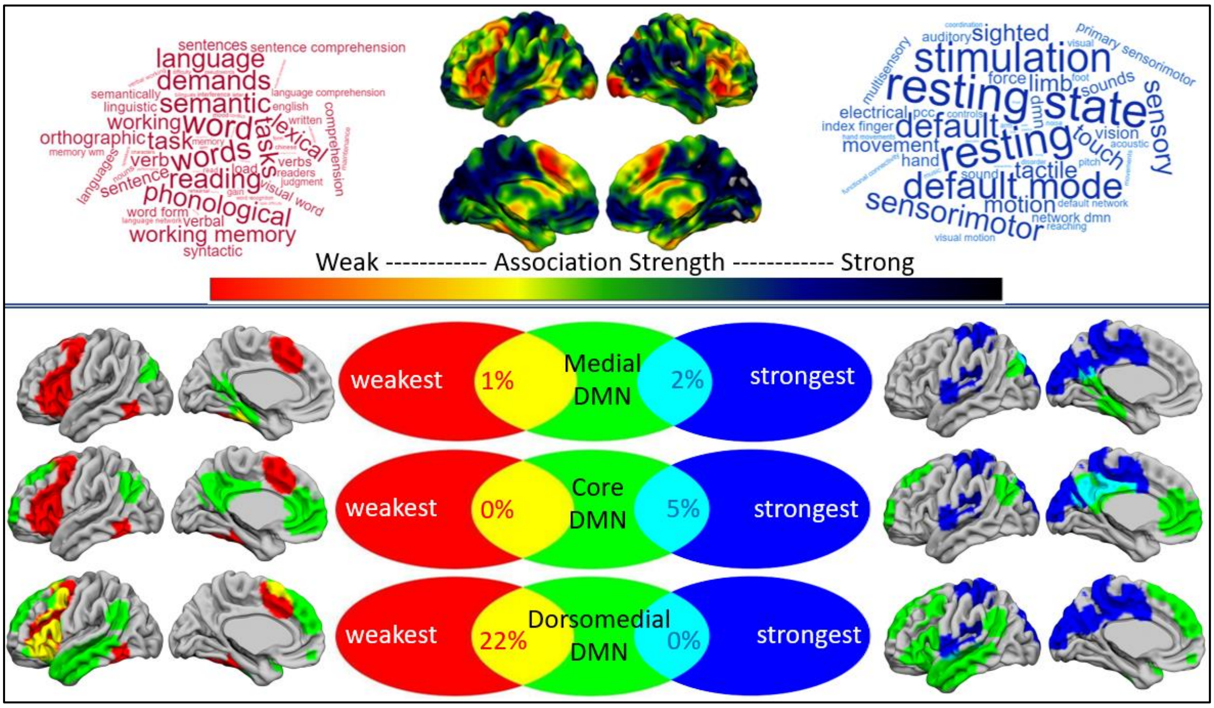


*Figure S3. Top panel: Unthresholded activation maps showing the continuous response associated with the parametric regressors. The word-clouds are derived from a Neurosynth meta-analysis of these maps. Bottom panel: Overlap of activation for parametric effects of strength of association, with the Default Mode Network (DMN) subsystems: medial (Yeo 15), core (Yeo 16) and dorsomedial (Yeo17). The Venn diagrams show the percentage of overlapping voxels for each end of the parametric effect with these established networks.*

<u>Correlations with Unusual Uses Task Performance</u>

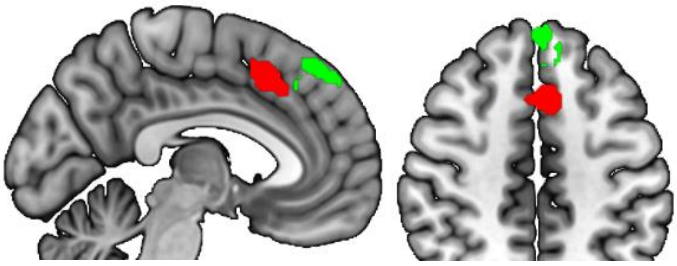

*Figure S4*

As a control analysis, we also confirmed that activation in a cluster adjacent to dmPFC (elicited by the episodic regressor) did not predict Unusual Uses performance. Activation in all three regressors (word2vec, uniqueness, episodic), across 4 ROIs, was used to predict performance on the Unusual Uses Task. The four ROIs were derived as follows: (1) Three ROIs from the uniqueness regressor: temporal fusiform gyrus, left IFG, dmPFC; one ROI derived from the episodic regressor adjacent to dmPFC (green in the figure above). This ROI comes from a larger swathe of activation in frontal cortex, but was restricted to the same extent as the dmPFC cluster, by excluding any activation that spread beyond medial frontal cortex, using the x co-ordinate from the uniqueness cluster (x = -12; see figure above; episodic cluster in green, uniqueness cluster in red). We conducted a series of control analyses to confirm that even with less predictors in the model, only activation for more unique response in dmPFC correlates with UUT (Tables S4-S6).

Dependent Variable: UUT

| Predictor (uniqueness activation) | Type III Sum of Squares | *df* | Mean Square | F | *p* |
|---|---|---|---|---|---|
| Intercept | 3.143 | 1 | 3.143 | 0.098 | 0.757 |
| temporal fusiform gyrus | 68.818 | 1 | 68.818 | 2.147 | 0.155 |
| left IFG | 75.363 | 1 | 75.363 | 2.351 | 0.137 |
| dmPFC | 260.435 | 1 | 260.435 | 8.124 | 0.008* |
| episodic medial frontal cortex | 41.99 | 1 | 41.99 | 1.31 | 0.263 |
| Error | 833.545 | 26 | 32.059 | | |

a R Squared = .283 (Adjusted R Squared = .173)

*Table S4. dmPFC is red in figure above, episodic medial prefrontal cortex is green in figure above.*

Dependent Variable: UUT

| Predictor (episodic activation) | Type III Sum of Squares | *df* | Mean Square | F | *p* |
|---|---|---|---|---|---|
| Intercept | 12.004 | 1 | 12.004 | 0.316 | 0.579 |
| temporal fusiform gyrus | 114.69 | 1 | 114.69 | 3.018 | 0.094 |
| left IFG | 125.574 | 1 | 125.574 | 3.305 | 0.081 |
| dmPFC | 30.82 | 1 | 30.82 | 0.811 | 0.376 |
| episodic medial frontal cortex | 44.399 | 1 | 44.399 | 1.169 | 0.29 |
| Error | 987.897 | 26 | 37.996 | | |

a R Squared = .150 (Adjusted R Squared = .020)

*Table S5. dmPFC is red in figure above, episodic medial prefrontal cortex is green in figure above.*

Dependent Variable: UUT

| Predictor (word2vec activation) | Type III Sum of Squares | *df* | Mean Square | F | *p* |
|---|---|---|---|---|---|
| Intercept | 7.032 | 1 | 7.032 | 0.163 | 0.689 |
| temporal fusiform gyrus | 9.488 | 1 | 9.488 | 0.221 | 0.643 |
| left IFG | 11.755 | 1 | 11.755 | 0.273 | 0.606 |
| dmPFC | 4.271 | 1 | 4.271 | 0.099 | 0.755 |
| episodic medial frontal cortex | 5.565 | 1 | 5.565 | 0.129 | 0.722 |
| Error | 1118.496 | 26 | 43.019 | | |

a R Squared = .038 (Adjusted R Squared = -.110)

*Table S6. dmPFC is red in figure above, episodic medial prefrontal cortex is green in figure above.*

Behavioural Data

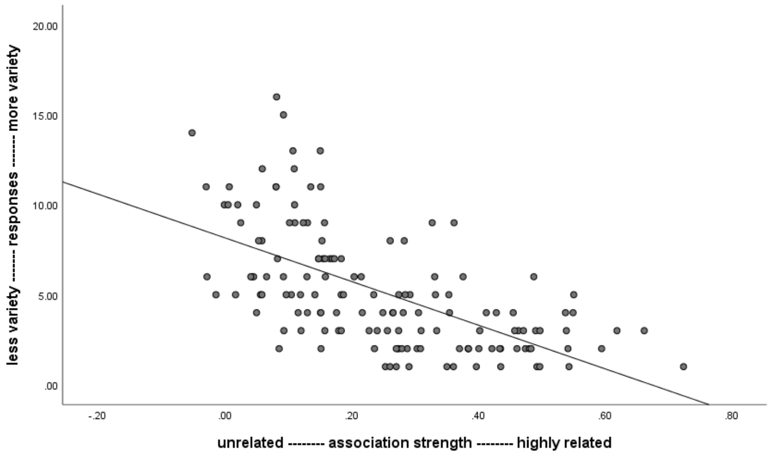


*Figure S5. Word pairs with the lowest word2vec scores (i.e., unrelated) had the greatest variety of responses across participants, while word-pairs with strong association strength tended to generate similar responses in different individuals (*Pearson r *= -.61,* p *< .001).*

There was a high degree of overlap between: participants' in-scanner ratings of the link they made and word2vec (*Pearson r* = .83, p < .001); word2vec and post-scan ratings (*Pearson r* = .82, p < .001); and in-scanner and post-scan ratings (*Pearson r* = .98, p < .001).

Participants were generally confident in their recall of the links they made in the scanner. 76% of recall was self-rated as highly confident (3 or 4 on a 0-4 scale), while only 12% of recall was rated as low confidence (0 or 1 on 0-4 scale). Confidence in recall does significantly correlate with associative strength (i.e., word2vec; *Pearson r* = .7, *p* < .001).

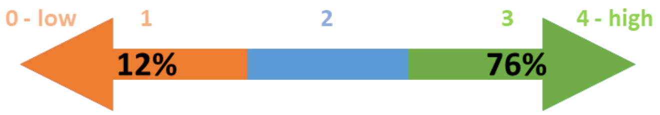


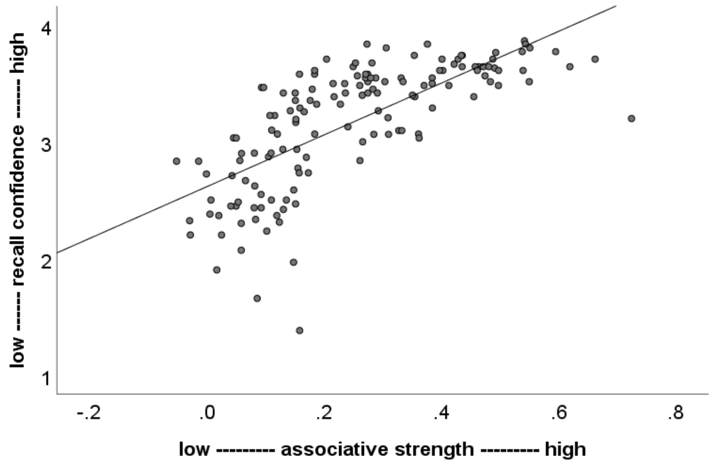

Even when removing the least confidently recalled items (i.e., those rated as 0 or 1), the correlation between uniqueness and associative strength (w2v) remained strong (*Pearson r* = .69, p < .001; the correlation for all trials is *Pearson r* = .72, p < .001).

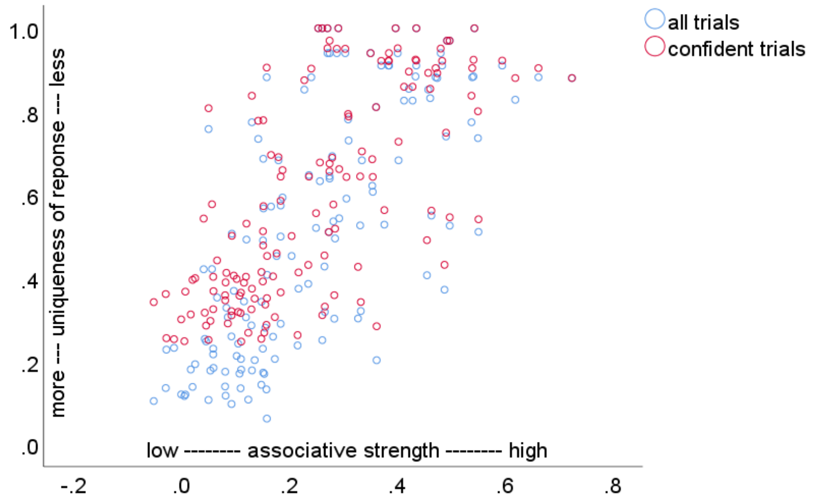
